# Geographic variation in sleep and metabolic function is associated with latitude and temperature

**DOI:** 10.1101/182790

**Authors:** Elizabeth B. Brown, Joshua Torres, Ryan A. Bennick, Valerie Rozzo, Arianna Kerbs, Justin R. DiAngelo, Alex C. Keene

## Abstract

Regulation of sleep and metabolic homeostasis are critical to an animal’s survival and under stringent evolutionary pressure. Animals display remarkable diversity in sleep and metabolic phenotypes; however, an understanding of the ecological forces that select for, and maintain, these phenotypic differences remain poorly understood. The fruit fly, *Drosophila melanogaster*, is a powerful model for investigating the genetic regulation of sleep and metabolic function, and screening in inbred fly lines has led to the identification of novel genetic regulators of sleep. Nevertheless, little is known about the contributions of naturally occurring genetic differences to sleep, metabolic phenotypes, and their relationship with geographic or environmental gradients. Here, we quantified sleep and metabolic phenotypes in 24 *D. melanogaster* populations collected from unique geographic localities. These studies reveal remarkable diversity in sleep, starvation resistance, and energy stores. We found that increased sleep duration is strongly associated with proximity to the equator and elevated average annual temperature, suggesting that environmental gradients strongly influence natural variation in sleep. Further, we found variation in metabolic regulation of sleep to be associated with free glucose levels, while starvation resistance associates with glycogen and triglyceride stores. Taken together, these findings reveal robust naturally occurring variation in sleep and metabolic traits in *D. melanogaster* and suggest that distance from the equator and median temperature is a significant evolutionary factor in sleep regulation and architecture.

## 1 | INTRODUCTION

Species display robust differences in homeostatically regulated behaviors including sleep, feeding and metabolic function, yet little is known about the ecological and functional relationship between these traits (Aulsebrook et al., 2016; Eban-Rothschild et al., 2017). In mammals, sleep duration ranges from ~2-18 hours per day, suggesting that the environment and evolutionary history potently affect sleep regulation (Capellini et al., 2008). While the ecological factors that drive differences in sleep remain largely unknown, a central hypothesis is that variation in sleep is linked to an animal’s metabolic and foraging needs. Supporting this notion, variation in sleep duration in *Drosophila* is associated with latitudinal cline (Svetec et al., 2015), raising the possibility that temperature and food availability also impact sleep need. Determining the relationship between sleep and metabolic function, and how evolution shapes these processes, is critical for understanding the function of sleep and basis for sleep differences between and within species.

The fruit fly *Drosophila melanogaster* presents a powerful model for investigating genetic interactions between sleep and metabolic processes (Erion et al., 2012; Yurgel et al., 2014). High throughput measurements of fly sleep and activity can be obtained using *Drosophila* Activity Monitors (DAMS), where infrared beam-breaks are indicative of fly movement (Pfeiffenberger et al., 2010a). Sleep is measured by five minutes of inactivity because it correlates with all other characteristics of sleep (Shaw et al., 2000). Flies acutely modulate their sleep in accordance with nutrient availability, and starvation potently inhibits sleep and initiates foraging, thereby providing a system to investigate the relationship between sleep and metabolic regulation (Keene et al., 2010; Lee and Park, 2004; Linford et al., 2012). Genetic evidence suggests sleep and metabolic function are highly conserved from flies to mammals at the molecular, pharmacological, and physiological levels (Allada and Siegel, 2008; Padmanabha and Baker, 2014), indicating that flies are an excellent system to examine the interactions between sleep and metabolic function.

While screens of inbred *Drosophila* lines have identified many regulators of sleep and metabolic regulation (Cirelli et al., 2005; Koh et al., 2008; Rogulja and Young, 2012), naturally occurring variation has also been leveraged to identify the genetic architecture regulating these processes (Harbison et al., 2009a, 2013). Quantitative genetic approaches in fully sequenced lines have provided insight into the genetic basis for resistance to environmental and physiological stressors including starvation resistance, and identified novel regulators of sleep (Harbison et al., 2004; Vieira et al., 2000). Further, experimental evolution and artificial selection approaches have revealed a relationship between sleep, feeding, and starvation resistance (Masek et al., 2014; Slocumb et al., 2015). For example, selection for short-sleeping flies results in reduced energy stores and sensitivity to starvation, while selecting for starvation resistance increases sleep duration (Masek et al., 2014; Seugnet et al., 2009). These studies have provided insight into the genetic and functional relationship between sleep and metabolic regulation, yet the ecological factors that shape the diversity of these traits in naturally occurring populations remain poorly understood.

Inspired by previous work investigating the relationship between geographic locality and sleep regulation (Svetec et al., 2015), we examined the relationship between sleep, metabolic function, and environmental localities. Here, we describe the analysis of sleep regulation, starvation resistance, the effects of starvation on sleep, and measurements of nutrient storage in *D. melanogaster* collected from geographically distinct localities that differ in latitude, longitude, temperature and altitude to determine the environmental and geographic factors that contribute to sleep regulation. We tested 24 populations of outbred *D. melanogaster* for these behavioral and physiological variables, providing insight into the relationship between these traits and their association with geographic locality. Our findings reveal highly significant variation in all traits measured, suggesting these traits are influenced by their environmental and evolutionary history.

## 2 | METHODS

### 2.1 | Drosophila maintenance

All populations were obtained from the *Drosophila* Species Stock Center (University of California, San Diego) with stock numbers provided in Table 1. Flies were reared and maintained on a 12:12 light-dark cycle in humidified incubators at 25°C and 65% humidity (Percival Scientific, Perry, IA). Unless otherwise noted, all flies were maintained and tested on standard cornmeal/agar medium.

**Table 1.** The localities of each *D. melanogaster* population and their corresponding geographic variables. Variables include latitude, longitude, altitude, and minimum temperature, maximum temperature, average temperature, and temperature range. For each population, its stock number and date of collection are also listed.

| Line | Stock | Locality | Latitude | Longitude | Altitude | Avg Temp |
| --- | --- | --- | --- | --- | --- | --- |
| 1 | 14021-0231.58 | Bermudas | 32°18'28.08"N | 64°45'1.80"W | 14.16 | 23.20 |
| 2 | 14021-0231.134 | American Samoa | 14°18'5.90"S | 170°41'46.25"W | 238.29 | 26.55 |
| 3 | 14021-0231.132 | Southwest Harbor, Maine | 44°16'47.37"N | 68°19'29.97"W | 11.35 | 6.58 |
| 4 | 14021-0231.59 | Bogota, Colombia | 4°42'39.56"N | 74° 4'19.53"W | 2581.03 | 20.51 |
| 5 | 14021-0231.69 | Athens, Greece | 37°59'1.71"N | 23°43'39.14"E | 71.08 | 16.95 |
| 6 | 14021-0231.64 | Koriba Dam, Zimbabwe | 16°31'19.58"S | 28°45'42.01"E | 664.14 | 23.34 |
| 7 | 14021-0231.131 | La Jolla, California | 32°49'58.12"N | 117°16'16.58"W | 51.89 | 16.49 |
| 8 | 14021-0231.136 | Fukushima, Japan | 37°45'39.00"N | 140°28'29.02"E | 68.24 | 11.72 |
| 9 | 14021-0231.67 | Pyrenees, Spain | 42°40'5.45"N | 1° 0'4.28"E | 2314.12 | 8.43 |
| 10 | 14021-0231.129 | Cebu, Philippines | 10°18'56.52"N | 123°53'7.57"E | 56.31 | 27.19 |
| 11 | 14021-0231.137 | Ogasawara Islands, Japan | 27° 4'30.19"N | 142°12'41.77"E | 223.45 | 24.38 |
| 12 | 14021-0231.61 | Blacksburg, Virginia | 37°13'46.46"N | 80°24'50.18"W | 634.67 | 11.58 |
| 13 | 14021-0231.23 | Crete, Greece | 35°14'24.42"N | 24°48'33.37"E | 1586.60 | 18.30 |
| 14 | 14021-0231.03 | Queensland, Australia | 20°55'3.27"S | 142°42'10.06"E | 321.11 | 25.09 |
| 15 | 14021-0231.43 | San Luis Potosi, Mexico | 22° 9'23.29"N | 100°59'7.95"W | 1877.06 | 18.85 |
| 16 | 14021-0231.15 | Bahia, Brazil | 12°34'47.06"S | 41°42'2.62"W | 715.73 | 22.13 |
| 17 | 14021-0231.130 | Queensferry, Scotland | 55°59'24.01"N | 3°23'56.56"W | 19.71 | 7.45 |
| 18 | 14021-0231.22 | Chiapas, Mexico | 16°45'24.95"N | 93° 7'45.25"W | 549.63 | 24.23 |
| 19 | 14021-0231.01 | Ica, Peru | 14° 4'31.66"S | 75°44'3.05"W | 399.25 | 19.25 |
| 20 | 14021-0231.35 | Monkey Hill, St. Kitts | 17°19'26.30"N | 62°43'30.14"W | 120.42 | 27.24 |
| 21 | 14021-0231.56 | Plainville, Connecticut | 41°40'32.68"N | 72°51'48.11"W | 52.86 | 9.74 |
| 22 | 14021-0231.62 | Capetown, South Africa | 33°55'29.53"S | 18°25'26.60"E | 52.57 | 16.35 |
| 23 | 14021-0231.68 | Isreal | 31° 2'45.78"N | 34°51'5.80"E | 287.34 | 19.93 |
| 24 | 14021-0231.66 | Madeira, Portugal | 32°45'38.55"N | 16°57'34.10"W | 1581.31 | 19.73 |

### 2.2 | Behavioral Analysis

Mated female flies aged 3-5 days were briefly anesthetized using CO_2_ and then individually placed into plastic tubes containing standard food. Flies were then acclimated to these conditions for at least 24hrs prior to testing. Fly activity was monitored using DAM2 *Drosophila* activity monitors (Trikinetics, Waltham, MA) as previously described (Hendricks et al., 2000; Shaw et al., 2000). The DAM system measures activity by monitoring the number of infrared beam crossings for each fly. These data were then used to calculate sleep-related traits by extracting immobility bouts of 5 minutes or more using the *Drosophila* Sleep Counting Macro (Pfeiffenberger et al., 2010b). Multiple variables of sleep were analyzed, including total sleep duration, waking activity, sleep bout number, and average sleep bout length as previously described (Pfeiffenberger et al., 2010b; Pitman et al., 2006). For experiments examining the effects of starvation on sleep, activity was recorded for one day on food prior to transferring flies into tubes containing 1% agar (Fisher Scientific, Hampton, NH). To calculate starvation-induced sleep suppression, the within-fly percentage change in sleep was calculated as ((% sleep starved-% sleep baseline)/% sleep baseline)*100 (Keene et al., 2010; Murakami et al., 2016). This calculation accounts for differences in baseline sleep and accurately reflects the response to starvation. For measurements of starvation resistance, flies were maintained on 1% agar until death. Time of death was manually determined as the last activity time point from the final recorded activity bout for each individual fly.

### 2.3 | Climate Data

Latitude, longitude, and altitude measurements for each locality were obtained from Google Earth (https://www.google.com/earth/). Latitude measurements were adjusted to represent the absolute distance from the equator. Longitude measurements to the east of the Prime Meridian were assigned a positive value, while measurements to the west were assigned a negative value. Since the altitude measurements were not normally distributed, the Log_10_ of altitude was used. Temperature data was obtained from Berkeley Earth (http://berkeleyearth.org/). Monthly temperature measurements were obtained from a 1°-by-1° latitude-longitude grid covering surface of the Earth. Average temperature was calculated as the average of the monthly temperature measurements.

### 2.4 | Triglyceride, glucose, and glycogen measurements

Protein, glucose, glycogen and triglyceride measurements were performed as previously described (Gingras et al., 2014; Sassu et al., 2012). Briefly, two headless female flies aged 3-5 days were homogenized in 50 mM Tris-HCl, pH 7.4, 140 mM NaCl, 0.1% Triton-X, and 1X protease inhibitor cocktail (Sigma Aldrich, St Louis, MO). Triglyceride levels were measured using the Infinity Triglycerides Kit (Fisher Scientific, Hampton, NH), while protein levels were measured using the Pierce BCA Protein Assay Kit (Fisher Scientific, Hampton, NH). Total glucose levels were determined using the Glucose Oxidase Reagent (Pointe Scientific, Canton, MI) in samples previously treated with 8mg/mL amyloglucosidase in 0.2M Sodium Citrate buffer, pH 5.0 (Boston BioProducts, Ashland, MA) for 2 hours. Glycogen levels were determined by measuring free glucose in samples not treated with amyloglucosidase and then subtracting the free glucose from total glucose concentration. For each sample, triglyceride, glycogen, and free glucose levels were standardized to the total protein content.

### 2.5 | Statistics

First, to assess the normality of each trait for each population we performed a Shapiro-Wilk test. For several traits, there was at least one non-normally distributed population, including: waking activity, bout length, % change in sleep, and measurements of glycogen levels. To assess variation among populations in traits where not all populations were normally distributed, we performed the nonparametric Wilcoxon Rank-Sum test. To assess variation among populations in traits in which all populations were normally distributed, we performed a one-way analysis of variance (ANOVA): Y = μ + Line + ε, where Line represents the fixed effect of population from a given locality and ε indicates error. In order to determine whether a given population differed from the global average, we performed a one-sample t-test and then adjusted for multiple testing using Bonferroni correction. A one-sample t-test was also performed to determine whether sleep responses as a result of starvation were significantly different from zero. Differences in survival upon starvation were assessed using the nonparametric log-rank test. Linear regressions were used to determine the relationship between a given trait and geographic variable as well as between two traits. For each correlation, the population average of each trait was used. All data was analyzed using JMP 12.0 software (SAS Institute Inc., Cary, NC).

## 3 | RESULTS

To determine the contribution of geographic variation on sleep and metabolic regulation, we obtained 24 populations of *D. melanogaster* from diverse localities throughout the world (Figure 1a). Outbred fly populations were selected based on their global distribution and variation in several geographic metrics, including latitude, longitude, altitude, and temperature (Table 1). To assess differences in sleep and metabolic function within individual flies, sleep was measured in 3-5-day-old female flies on food. Following 24hrs of sleep acquisition, flies were transferred to starvation tubes containing agar alone and maintained on this substrate to measure starvation-induced changes in sleep and starvation resistance (Figure 1b), providing multiple metrics of sleep and metabolic function within an individual animal.

**FIGURE 1.**
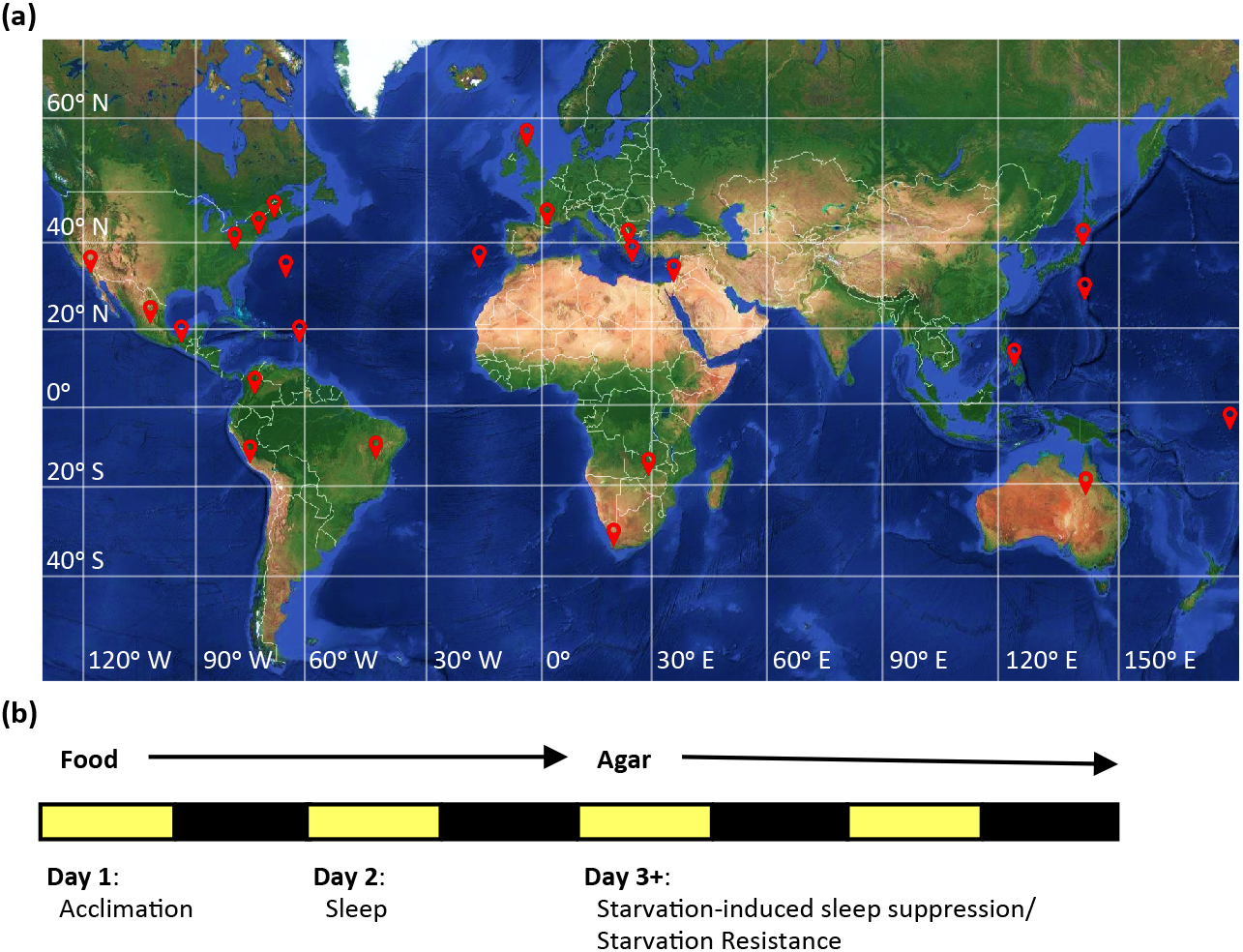
Localities of *D. melanogaster* populations and experimental design. (a) World map displaying the localities (red pins) of each the 24 populations. Horizontal lines indicate latitude and vertical lines indicate longitude. (b) Schematic of experimental protocol used to assess sleep- and metabolic-related traits.

### 3.1 | Variation in sleep traits

Quantification of sleep duration in fed flies revealed remarkable diversity in sleep duration and architecture between fly populations (Figure 2). The average sleep duration of all 24 lines tested for 24 hrs on food was 855 min. The shortest sleeping populations include flies from Fukushima, Japan and Israel, sleeping on average 413 and 570 min, respectively (Figure 2A). In addition, several long-sleeping populations were identified, including flies from Bogota, Columbia (1100 min) and Bermuda (1117 min) (Figure 2a). The differences between short- and long-sleeping flies were present during both the day and night, thereby suggesting that the observed phenotypes are not the result of altered circadian regulation (Figure 2b).

**FIGURE 2.**
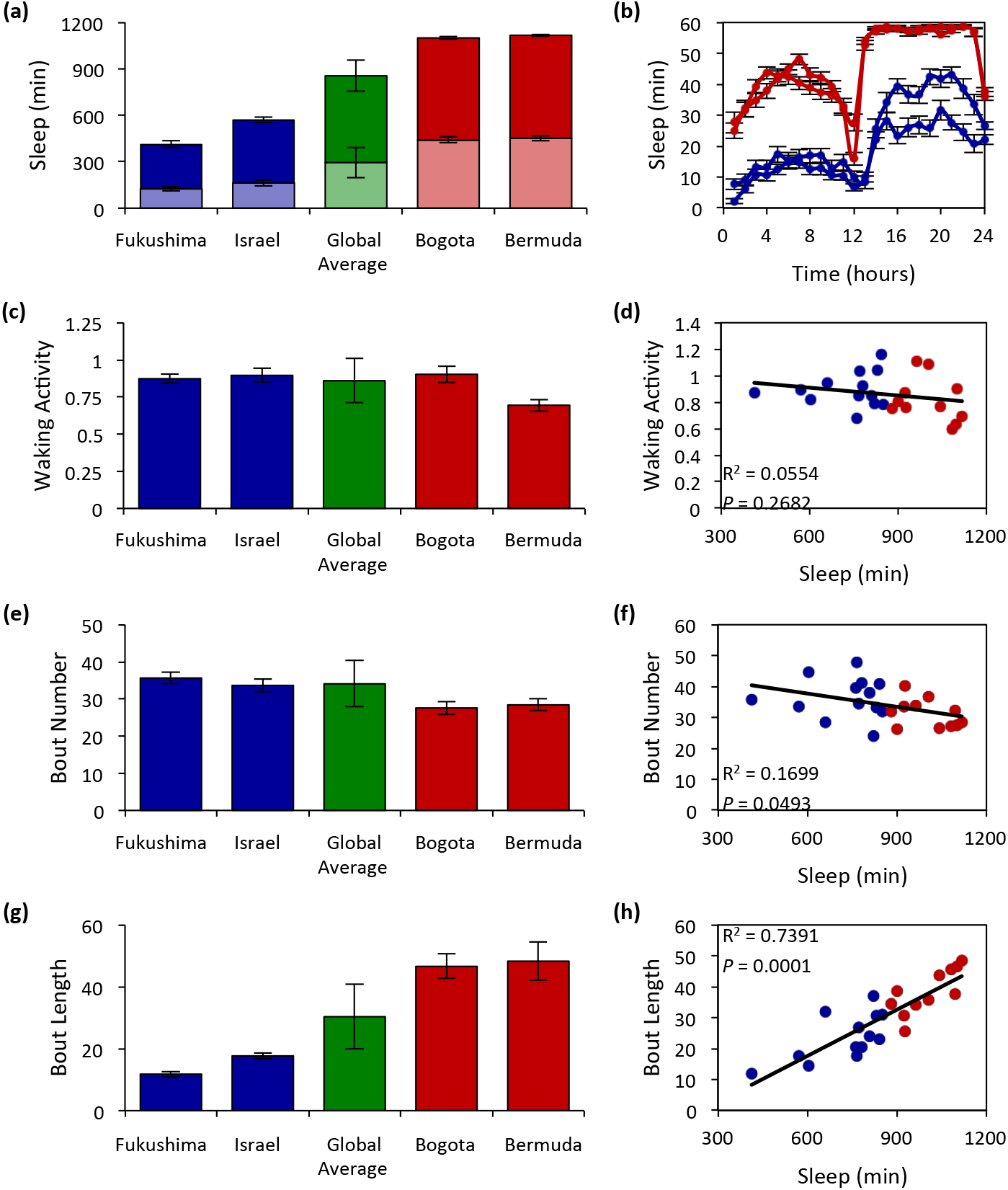
Measurements of sleep-related traits reveal significant variation among *D. melanogaster* populations. (a) Daytime (light bars) and nighttime (dark bars) sleep duration for the two shortest and longest sleeping populations relative to the global average. There is significant variation in both daytime and nighttime sleep duration among the populations tested (daytime: ANOVA: *F*_23,811_ = 23.2805, *P* < 0.0001; nighttime: ANOVA: *F*_23,811_ = 39.2778, *P* < 0.0001). (b) Sleep profiles over a 24hr period of the two shortest and longest sleeping populations. (c) Measurements of waking activity for the two shortest and longest sleeping populations relative to the global average. There is significant variation in waking activity among the populations tested (Wilcoxon rank-sum test: *Χ*^2 =^ 240.7045, *P* < 0.0001). (d) There is no correlation between sleep duration and waking activity. (e) Measurements of bout number for the two shortest and longest sleeping populations relative to the global average. There is significant variation in bout number among the populations tested (ANOVA: *F*_23,811_ = 11.7775, *P* < 0.0001). (f) There is a moderate, but significant correlation between sleep duration and bout number. (g) Measurements of bout length for the two shortest and longest sleeping populations relative to the global average. There is significant variation in bout length (Wilcoxon rank-sum test: *Χ*^2 =^ 340.777, *P* < 0.0001) among the populations tested. (h) There is a significant correlation between sleep duration and bout length. For each population, data shown is mean ± SE. For global averages, data shown is mean ± SD. N = 30-40. Blue dots represent populations whose sleep duration is below the global average, while red dots represent populations above the global average. For measurements of sleep-related traits for all localities see Appendix 1.

It is possible that differences in sleep duration are reflective of lethargy or hyperactivity rather than sleep per se. To investigate this, we measured the relationship between sleep duration and waking activity. We observed no difference in waking activity between the two shortest sleeping populations and the global average, while a moderate reduction was observed in one of the long sleeping populations (Figure 2c). A regression analysis of the 24 populations tested revealed a lack of correlation between sleep duration and waking activity (Figure 2d), suggesting that these traits are independently regulated and that the differences in sleep across geographic localities are not due to hyperactivity or lethargy.

Sleep duration is composed of individual sleep bouts and may be increased by increasing the number of bouts or the length of each individual sleep bout within a given day (Pfeiffenberger et al., 2010b). To differentiate between these possibilities, we calculated the average number of bouts for each population. Overall, the average number of bouts ranged from 24-48 per 24 hours, and there was a moderate but significant correlation between bout number and sleep duration, suggesting that sleep initiation only mildly contributes to variation in sleep duration (Figure 2e,f). In addition, the length of individual sleep bouts were shorter in the Fukushima, Japan and Israel populations, and longer in the Bogota, Columbia and Bermuda populations compared to the global average (Figure 2g). Furthermore, a regression analysis revealed a highly significant correlation between sleep duration and bout length (Figure 2h). Taken together, these findings suggest that the diversity in sleep duration is primarily conferred by increasing the length of individual sleep bouts.

A previous study examining *D. melanogaster* collected from five North American localities suggested that sleep is increased in equatorial regions (Svetec et al., 2015). To determine how sleep and activity relate to variation in geographic and environmental locality, we performed linear regressions on four geographic variables including latitude, longitude, altitude, and temperature. This analysis revealed a significant relationship between sleep duration and both latitude and temperature, while no relationship was observed between sleep duration and longitude or altitude (Figure 3, Table 2). Specifically, we observed a significant negative correlation between sleep duration and latitude (Figure 3a), while a significant positive correlation was observed between sleep duration and temperature (Figure 3b). We observed a significant positive correlation between waking activity and latitude, while a significant negative correlation was observed between waking activity and temperature. Taken together, these results suggest that close proximity to the equator and increased temperature are associated with increased sleep duration and reduced waking activity.

**FIGURE 3.**
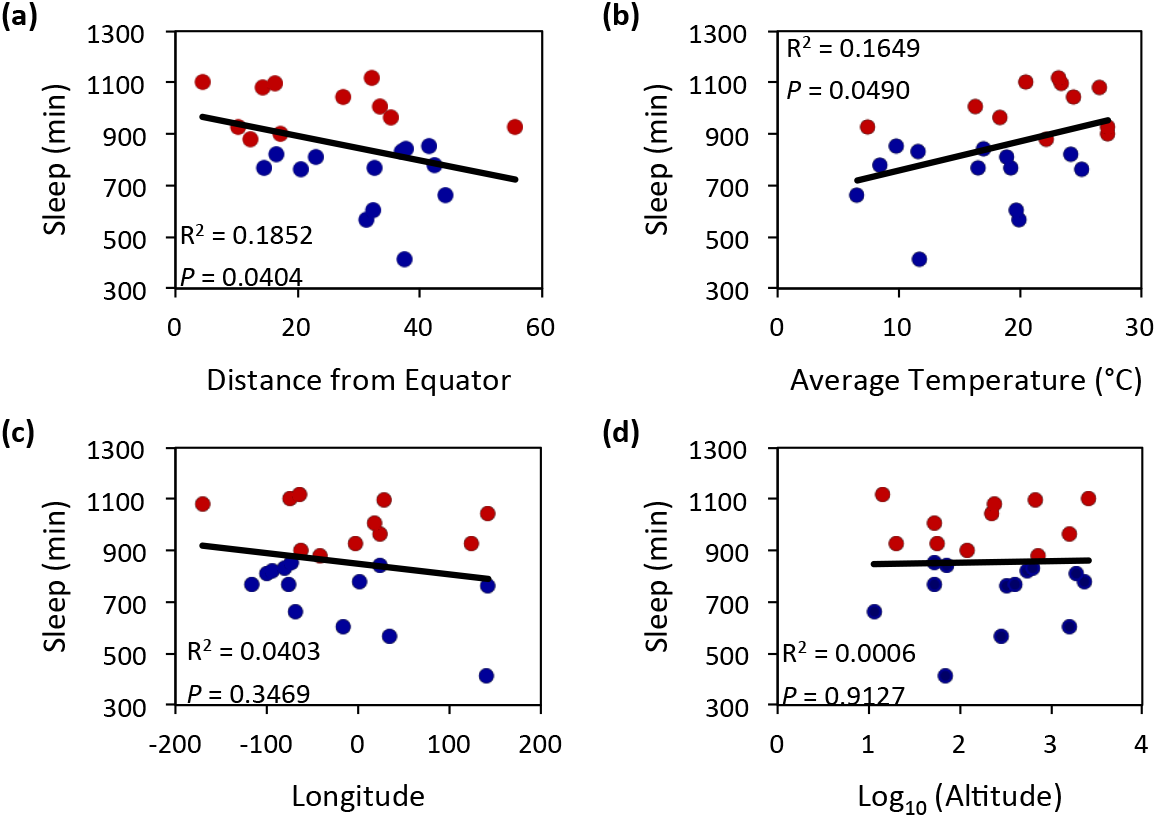
Linear regression analyses highlight the relationship between sleep duration and geographic variables. There is a significant relationship between (a) sleep duration and distance from the equator as well as (b) sleep duration and temperature. There is no relationship between (c) sleep duration and longitude nor between (d) sleep duration and altitude. Blue dots represent populations whose sleep duration is below the global average, while red dots represent populations above the global average. Statistical analyses are reported in Table 2.

**Table 2.** Results of linear regression analyses predicting sleep traits from geographic variables. Significant correlations are shown in bold.

| Trait: | Geographic Variable: | F-Value | P-Value | R<sup>2</sup> |
| --- | --- | --- | --- | --- |
| Sleep Duration (Day) | Latitude | $F_{1,22} = 2.6100$ | 0.1211 | 0.1105 |
| | Longitude | $F_{1,22} = 0.0008$ | 0.9781 | 0.0000 |
| | Altitude | $F_{1,22} = 0.0574$ | 0.8129 | 0.0026 |
| | Temperature | $F_{1,22} = 3.0298$ | 0.0957 | 0.0034 |
| Sleep Duration (Night) | Latitude | $F_{1,22} = 5.1586$ | **0.0338** | 0.1972 |
| | Longitude | $F_{1,22} = 3.4109$ | 0.0783 | 0.1342 |
| | Altitude | $F_{1,22} = 0.0015$ | 0.9698 | 0.0000 |
| | Temperature | $F_{1,22} = 3.8453$ | 0.0627 | 0.1489 |
| Sleep Duration (24hrs) | Latitude | $F_{1,22} = 4.7732$ | **0.0404** | 0.1852 |
| | Longitude | $F_{1,22} = 0.9239$ | 0.3469 | 0.0403 |
| | Altitude | $F_{1,22} = 0.0123$ | 0.9127 | 0.0006 |
| | Temperature | $F_{1,22} = 4.3434$ | **0.0490** | 0.1649 |
| Activity | Latitude | $F_{1,22} = 6.5475$ | **0.0179** | 0.2294 |
| | Longitude | $F_{1,22} = 0.0020$ | 0.9647 | 0.0000 |
| | Altitude | $F_{1,22} = 0.0247$ | 0.8765 | 0.0011 |
| | Temperature | $F_{1,22} = 8.5920$ | **0.0077** | 0.2809 |
| Bout Number | Latitude | $F_{1,22} = 0.3478$ | 0.5614 | 0.0156 |
| | Longitude | $F_{1,22} = 1.7009$ | 0.2056 | 0.0718 |
| | Altitude | $F_{1,22} = 0.8147$ | 0.3765 | 0.0357 |
| | Temperature | $F_{1,22} = 0.7955$ | 0.3821 | 0.0349 |
| Bout Length | Latitude | $F_{1,22} = 2.0709$ | 0.1642 | 0.0860 |
| | Longitude | $F_{1,22} = 2.9394$ | 0.1005 | 0.1179 |
| | Altitude | $F_{1,22} = 0.2444$ | 0.6260 | 0.0110 |
| | Temperature | $F_{1,22} = 2.9562$ | 0.0996 | 0.1185 |

### 3.2 | Variation in the metabolic regulation of sleep

Flies, like mammals, suppress sleep when starved, presumably to initiate a foraging response (Danguir and Nicolaidis, 1979; Keene et al., 2010). This phenotype has been extensively investigated in inbred fly lines, however it has not been studied in outbred populations of *Drosophila* (Yurgel et al., 2014). To determine how sleep is modulated by nutrient deprivation, flies were starved following 24-hour sleep recordings on food by transferring flies to tubes containing agar alone and the change in sleep between the two housing conditions was then determined. Fourteen lines significantly suppressed sleep in response to starvation, eight lines exhibited no change in sleep, and two lines significantly increased sleep (Appendix S2). Flies from Cape Town, South Africa (44%) and Koriba Dam, Zimbabwe (42%) displayed the greatest sleep suppression, while flies from Plainville, Connecticut (10%) and Madeira, Spain (39%) showed the greatest increase (Figure 4a). There was no correlation between sleep duration on food and starvation-induced sleep suppression (Figure 4b), indicating that sleep duration and changes in sleep resulting from starvation are independently regulated.

**FIGURE 4.**
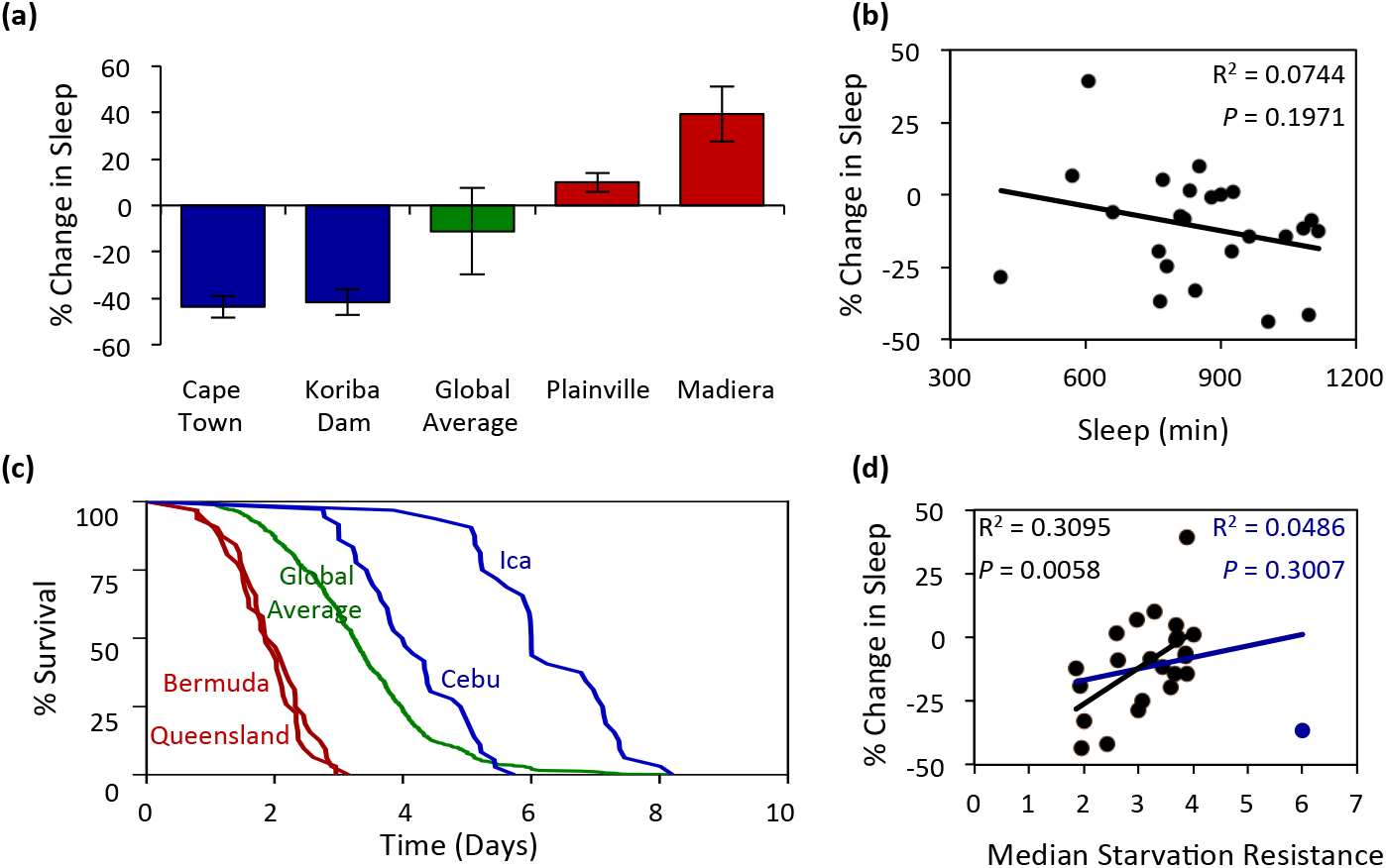
The effect of starvation on sleep and longevity in *D. melanogaster* populations. (a) There is significant variation in starvation-induced changes in sleep duration (Wilcoxon rank-sum test: *Χ*^2=^ 185.9639, *P* < 0.0001). The change in sleep for the two shortest and longest sleeping populations are shown relative to the global average. For each population, data shown is mean ± SE. For global average, data shown is mean ± SD. N = 30-40. (b) There is no relationship between sleep duration and starvation-induced changes in sleep. (c) There is significant variation in measurements of starvation resistance (Log-rank test: *Χ*^2 =^ 811.1161, *P* < 0.0001). Survivorship curves for the two least and most starvation resistant populations are shown relative to the global average. N ¯ 30-40. (d) The relationship between starvation resistance and starvation-induced changes in sleep. The blue regression line includes Ica, Peru, the outlier population (blue dot), while the black regression lines does not. For measurements of starvation-induced sleep suppression and starvation resistance for all localities see Appendix 1.

It is possible that sleep suppression in response to starvation represents a specific sensitivity to sleep regulation, or provides a more general indicator of starvation tolerance. To measure starvation resistance, we maintained flies on agar in activity monitors and measured time of death. The mean starvation resistance for all 24 lines tested was 3.26 days. The least resistant populations include flies from Bermuda and Queensland, Australia, living on average 1.86 and 1.93 days, respectively. The most resistant populations include flies from Cebu, Philippines, living 4.01 days and Ica, Peru, living 6.00 days (Figure 4c). We observed no correlation between starvation-induced sleep suppression and starvation resistance when all flies were included in the regression (blue line Figure 4d). Given that flies from Ica, Peru live approximately 2 days longer than the next longest-living population and over 3 standard deviations above the global average, we next performed a linear regression in the absence of this outlier locality. We found a strong positive correlation between starvation-induced sleep suppression and starvation resistance (black line: Figure 4d), suggesting that sleep duration and changes in sleep resulting from starvation are not independently regulated. To determine whether starvation-induced sleep suppression and/or starvation resistance is associated with variation in geographic variables, we performed linear regression analyses between these traits and the geographic variables. These analyses revealed no correlation between neither starvation-induced sleep suppression nor starvation resistance with any geographic variables measured (Table 3). Taken together, these findings demonstrate dramatic variation in starvation resistance and the metabolic regulation of sleep in different fly populations, indicating that the metabolic regulation of sleep is coregulated with starvation resistance.

**Table 3.** Results of linear regression analyses predicting starvation-induced sleep suppression and starvation resistance from geographic variables.

| Trait: | Geographic Variable: | F-Value | P-Value | R<sup>2</sup> |
| --- | --- | --- | --- | --- |
| % Change in Sleep | Latitude | $F_{1,22} = 0.0000$ | 0.9971 | 0.0000 |
| | Longitude | $F_{1,22} = 1.5103$ | 0.2321 | 0.0642 |
| | Altitude | $F_{1,22} = 0.2861$ | 0.5981 | 0.0128 |
| | Temperature | $F_{1,22} = 0.1061$ | 0.7477 | 0.0048 |
| Starvation Resistance | Latitude | $F_{1,22} = 0.5774$ | 0.4554 | 0.02558 |
| | Longitude | $F_{1,22} = 0.9688$ | 0.3357 | 0.0422 |
| | Altitude | $F_{1,22} = 0.3075$ | 0.5848 | 0.0138 |
| | Temperature | $F_{1,22} = 0.0000$ | 0.9968 | 0.0000 |

### 3.3 | Variation in nutrient storage

Energy stores and circulating nutrients potently affect both sleep and starvation resistance. To investigate the relationship between energy stores and these processes, we measured triglyceride levels, glycogen levels, and free glucose across all 24 populations. We identified at least a 4-fold difference in triglyceride, glycogen, and free glucose levels in the populations tested (Figure 5a-c). Across all three measurements, energy stores were lowest in the Queensland, Australia population. A linear regression analysis of the relationship between energy levels and starvation-induced sleep suppression revealed no significant correlation between starvation-induced changes in sleep and triglyceride or glycogen levels (Figure 5d,e). However, a significant correlation was observed between starvation-induced regulation of sleep and free glucose, suggesting that flies with lower levels of free glucose during the fed state display a greater reduction in sleep upon starvation (Figure 5f). Conversely, increased starvation resistance correlated with elevated levels of triglycerides and glycogen, but not free glucose (Figure 5g-i), indicating elevated energy stores in fed flies are indicators of starvation resistance. Taken together, these findings suggest that energy storage molecules (glycogen and triglycerides) and free glucose have distinct effects on the metabolic regulation of sleep and starvation resistance.

**FIGURE 5.**
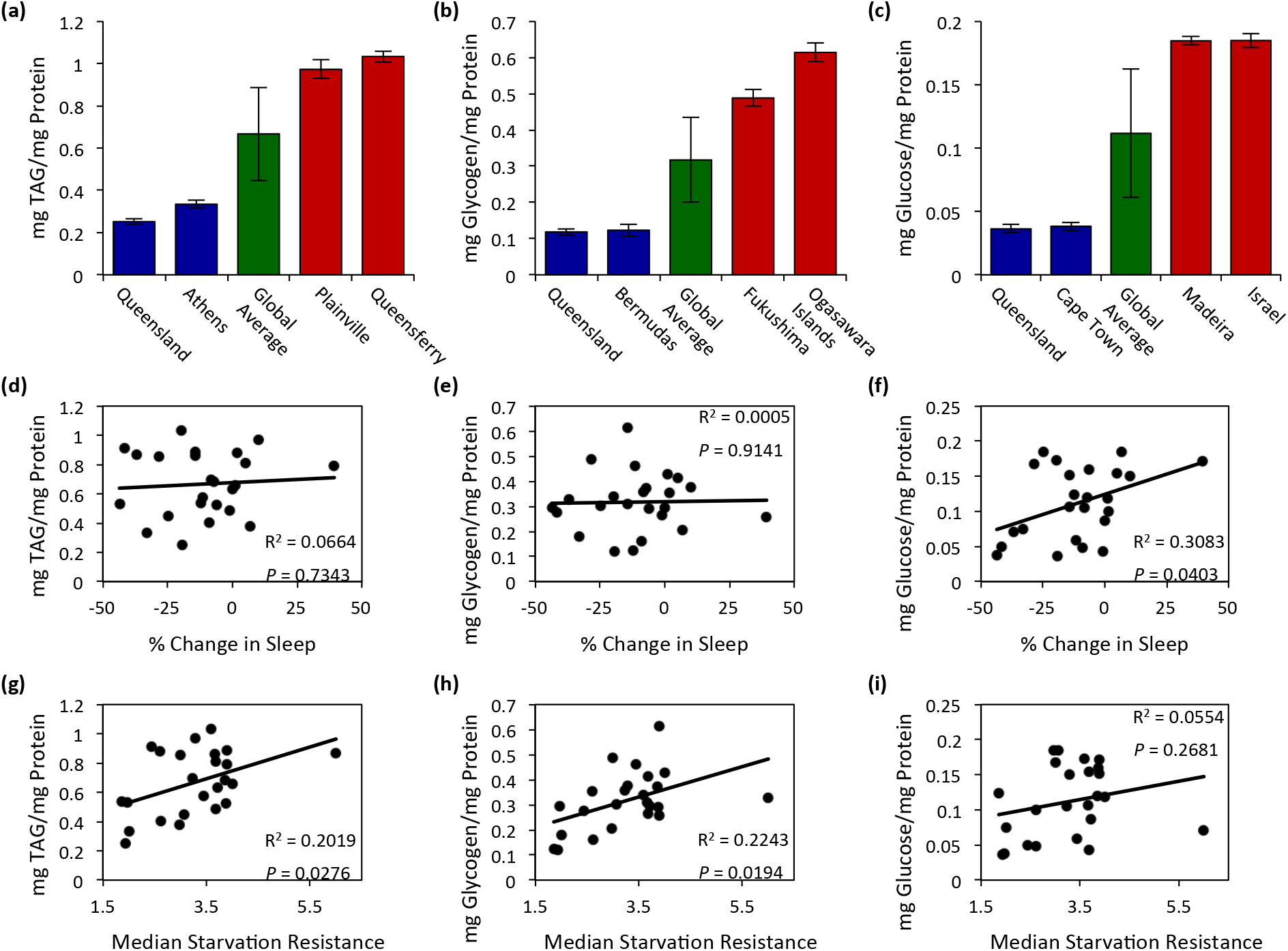
Energy storage measurements of *D. melanogaster* populations. (a) There is significant variation in triglycerides (a; ANOVA: *F*_23,309_ = 34.8198, *P* < 0.0001, N = 915), glycogen (b; Wilcoxon rank-sum test: *Χ*^2 =^ 202.5611, *P* < 0.0001, N = 10-15), and free glucose levels (c; ANOVA: *F*_23,310_ = 79.1969, *P* < 0.0001, N = 9-15). For each trait, the two populations with the lowest and highest measurements are shown relative to the global average. For each population, data shown is mean ± SE. For global average, data shown is mean ± SD. (d-f) The relationships between starvation-induced sleep suppression and energy storage measurements. There is no correlation between starvation-induced sleep suppression and (d) triglyceride or (e) glycogen levels. However, starvation-induced sleep suppression was significantly correlated with (f) free glucose levels. (g-i) The relationships between starvation resistance and energy storage measurements. There is a significant correlation between starvation resistance and (g) triglyceride and (h) glycogen levels, while there is no correlation between starvation resistance and (i) free glucose levels. For measurements of nutrient stores for all localities see Appendix 1.

To determine the relationship between energy stores and free glucose with geographic variables, we again performed linear regression analyses between these traits and the geographic variables. These analyses revealed no correlation between triglyceride or glycogen levels and any geographic variables measured (Table 4). Conversely, there was a correlation between free glucose levels and geographic variables (Figure 6). We found that free glucose levels were significantly correlated with increased distance from the equator and decreased mean annual temperatures (Figure 6a,b; Table 4). Therefore, shared environmental pressures appear to contribute to the regulation of sleep duration and free glucose.

**FIGURE 6.**
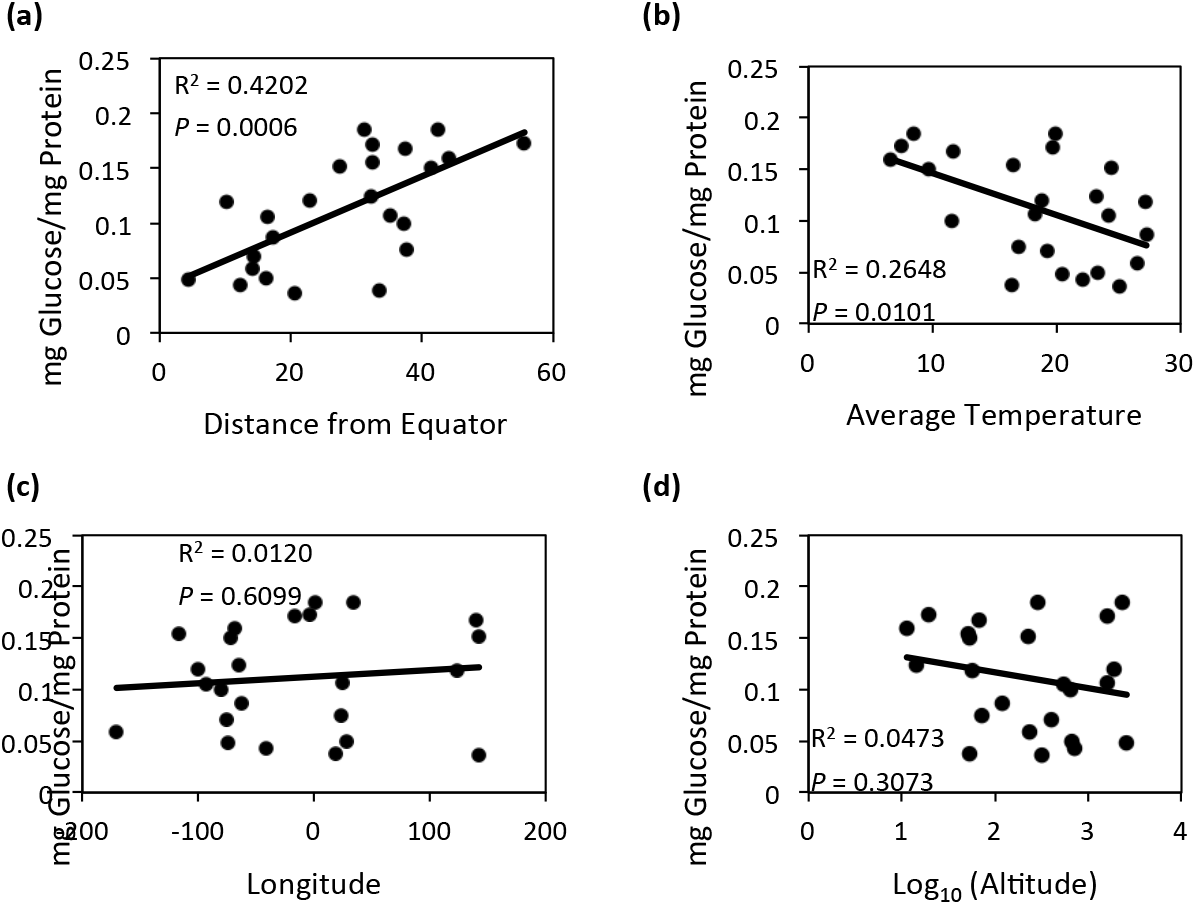
Linear regression analyses highlight the relationship between free glucose and geographic variables. There is a significant positive relationship between (a) free glucose levels and distance from the equator, while there is a significant negative relationship between (b) free glucose and temperature. There is no relationship between free (c) glucose levels and longitude nor between (d) free glucose and altitude. Statistical analyses are reported in Table 4.

**Table 4.**
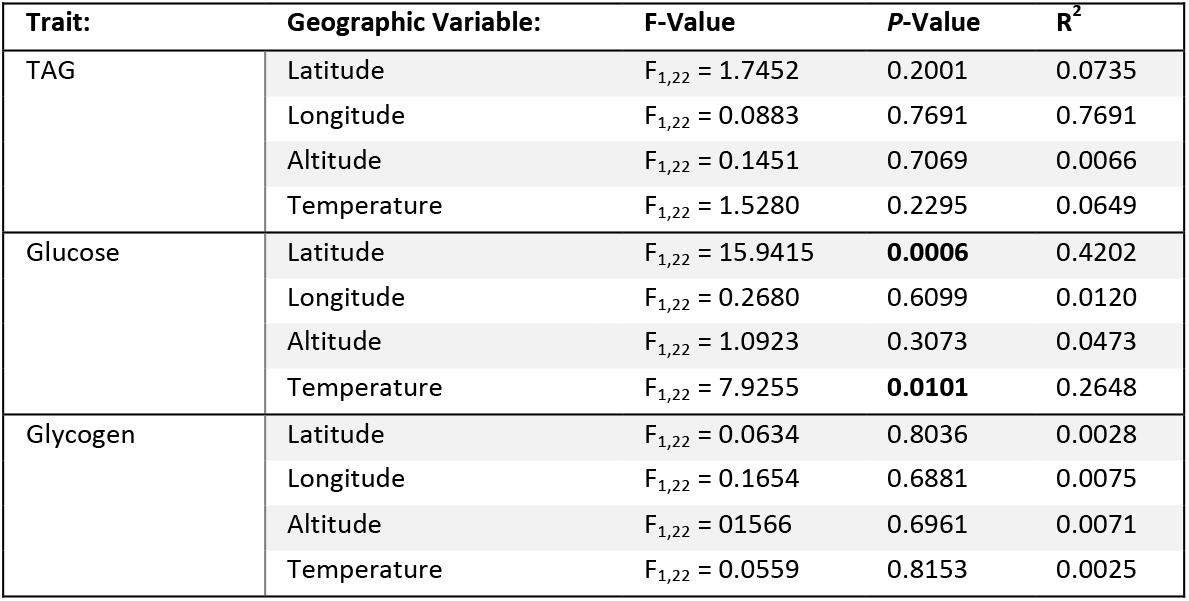
Results of linear regression analyses predicting energy storage traits from geographic variables. Significant correlations are shown in bold.

| Trait: | Geographic Variable: | F-Value | P-Value | R<sup>2</sup> |
| --- | --- | --- | --- | --- |
| TAG | Latitude | $F_{1,22} = 1.7452$ | 0.2001 | 0.0735 |
| | Longitude | $F_{1,22} = 0.0883$ | 0.7691 | 0.7691 |
| | Altitude | $F_{1,22} = 0.1451$ | 0.7069 | 0.0066 |
| | Temperature | $F_{1,22} = 1.5280$ | 0.2295 | 0.0649 |
| Glucose | Latitude | $F_{1,22} = 15.9415$ | **0.0006** | 0.4202 |
| | Longitude | $F_{1,22} = 0.2680$ | 0.6099 | 0.0120 |
| | Altitude | $F_{1,22} = 1.0923$ | 0.3073 | 0.0473 |
| | Temperature | $F_{1,22} = 7.9255$ | **0.0101** | 0.2648 |
| Glycogen | Latitude | $F_{1,22} = 0.0634$ | 0.8036 | 0.0028 |
| | Longitude | $F_{1,22} = 0.1654$ | 0.6881 | 0.0075 |
| | Altitude | $F_{1,22} = 0.1566$ | 0.6961 | 0.0071 |
| | Temperature | $F_{1,22} = 0.0559$ | 0.8153 | 0.0025 |

## 4 | DISCUSSION

The expansive radiation of *Drosophila melanogaster* provides an excellent opportunity to examine the interrelationship between natural variation in complex behaviors, physiological traits, and the environmental factors that may act as selective forces to shape such variation. Here we examined natural variation in sleep and metabolic function, as well as their relationship with each other, and with several geographic and environmental gradients, by investigating 24 *D. melanogaster* populations from localities across the globe. The vast majority of sleep studies in *Drosophila* have used inbred or isogenic *D. melanogaster* strains. Inbred populations of laboratory *Drosophila* strains typically sleep ~600-800 min daily (Zimmerman et al., 2012), while the sleep duration for strains in this study ranged from 413-1117 minutes. Similarly, starvation resistance in inbred strains is ~36-49 hrs, while flies in this study survived up to 6 days (Gáliková et al., 2015; Mattaliano et al., 2007). Therefore, naturally occurring variability encompasses the sleep and starvation resistant phenotypes observed in the most commonly used laboratory strains. This remarkable variability in natural populations of *Drosophila* is suggestive of maintenance of genetic and phenotypic variation associated with geographically independent populations of *Drosophila.* The extreme phenotypes observed in natural populations of *Drosophila* suggest differences between populations could be used to uncover novel genetic architecture associated with natural variation in sleep and metabolic function.

Sleep is regulated by complex genetic architecture and is highly influenced by genetic variation. While many genes have been identified using mutagenesis approaches, much less is known about the modulation of sleep via naturally occurring genetic variation. In humans, sleep need is estimated to range from as 7-9 hours, and single alleles have been identified that robustly influence sleep (Allebrandt et al., 2013; Shi et al., 2017). Genome-wide association studies have identified loci associated with sleep variability, but assessing the contributions of loci to sleep variation is difficult (Kalmbach et al., 2017). Sequenced inbred *Drosophila* lines derived from a wild-caught population have been previously characterized for variation in sleep, revealing numerous differentially expressed genes and molecular polymorphisms that associate with sleep duration and architecture (Harbison et al., 2009b, 2013). The identification of differences in sleep and metabolic phenotypes from geographically diverse regions presents a complementary approach to investigate the genetic architecture underlying variation in sleep regulation.

Clinal variation has been observed in diverse traits including body size, fecundity, and temperature resistance, but much less is known about the relationship between behavior and clinality (for review see (Adrion et al., 2015)). Here, we identify a relationship between sleep duration and two environmental gradients: distance from the equator and average temperature. Specifically, we found that increased sleep duration is associated with proximity to the equator and increased average temperature. These findings are in agreement with a previous study using five North American populations in which they found clinal variation in nighttime bout length and sunrise anticipation (Svetec et al., 2015). This study revealed that the clinal relationship between latitude and sleep were observed during the nighttime only, suggesting that ecological variables have differential effects on daytime and nighttime sleep (Svetec et al., 2015). Similarly, we observed no relationship between latitude and daytime sleep, further supporting the notion that clinal variation in sleep duration is nighttime specific. Unsurprisingly, there are many additional traits associated with latitudinal clines. Sunrise anticipation, where flies become active prior to the onset of the light cycle, was also associated with latitude (Svetec et al., 2015). Both morning anticipation and sleep duration are regulated by the circadian clock in *Drosophila.* Therefore, it is possible that alterations in circadian neural circuitry or the molecular machinery governing the circadian clock contributes to cline-associated differences in sleep among *Drosophila* populations.

Flies robustly suppress sleep and increase activity in response to starvation, and this is mediated by both chemosensory and hormonal factors (Keene et al., 2010; Lee and Park, 2004; Linford et al., 2012; Murakami et al., 2016). While this phenotype has been reported in diverse genetic backgrounds of *D. melanogaster* (McDonald and Keene, 2010), we found that 10 of the 24 populations tested do not suppress sleep when starved. It is possible that this is associated with a delayed response to starvation, and analysis of sleep during later periods of starvation will identify starvation-induced sleep suppression. Surprisingly, we identified two populations that increased sleep during starvation, revealing opposing responses to starvation. Although enhanced sleep during starvation has not been reported in *Drosophila*, we have previously identified increased sleep during starvation in the Mexican blind cavefish, and many animals enter hibernation during winter seasons when food availability is scarce (Jaggard et al., 2017; Schmidt, 2014). Therefore, there are likely multiple strategies that are implemented in response to food shortage, including induction of foraging behavior and consequently sleep suppression, or increasing sleep to conserve energy. A better understanding of how environmental factors shape the evolution of these opposing strategies are of particular interest.

In this study, we did not find an association between starvation resistance or starvation-induced sleep suppression with any environmental gradient, raising the possibility that (1) variation in these traits may be due to an environmental gradient not measured in this study, or (2) these traits do not correlate with clinal variables. Nevertheless, in the case of starvation resistance, several previous studies have also failed to find a relationship with latitudinal cline (Goenaga et al., 2010; Hoffmann et al., 2005; Robinson et al., 2000), while others reported negative latitudinal clines (Arthur et al., 2008; Karan and Parkash, 1998; Karan et al., 1998). To our knowledge, no similar investigation has been previously performed on the relationship between environmental gradients and starvation-induced sleep suppression. This suggests that clinal variation in starvation resistance may vary depending on the localities of the populations tested. Although we did not identify clinal variation in starvation resistance or starvation-induced sleep suppression, our analysis does not take into account additional variables such as ultraviolet light intensity, seasonality, day length, or other factors that may potently affect food availability and influence selection. It is also possible that there are environmental factors specific to individual localities that shape the behavioral and metabolic responses we observed. One population in which this may indeed be the case is *Drosophila* from Ica, Peru, the most starvation resistant population in our analysis. The city of Ica borders the Atacama Desert, which is classified as a hot desert climate according to the Köppen-Geiger climate classification system (Peel et al., 2007). We posit that the dry, arid climate of this region would select for *Drosophila* with an increased resistance to starvation, as this is the case with several *Drosophila* species that are found in more xeric habitats (Matzkin and Markow, 2009). Nevertheless, our findings that latitudinal cline and temperature are associated with differences in sleep duration and activity across continents and geographic conditions suggest a strong relationship between latitude and sleep regulation.

Upon investigation into the relationship between sleep regulation and energy stores, we found that populations with lower levels of free glucose display a greater increase in sleep suppression after starvation. Limiting glucose utilization pharmacologically suppresses sleep, presumably mimicking the starvation state. (Murakami et al 2016). These findings suggest natural variation in free glucose impact how flies modulate sleep in accordance with nutrient shortage. We also found a positive correlation between energy storage measurements (glycogen and triglyceride levels) and starvation resistance. Previous work has shown that artificial selection for increased starvation resistance results in a correlated increase in energy stores (Slocumb et al., 2015), raising the possibility that increased energy stores are an adaption to areas with limited or sporadic food availability. Of these metabolic traits, we only observed free glucose associates with equatorial proximity. Interestingly, allozymes of glucose-6-phosphate dehydrogenase (G6PD), an enzyme involved in the breakdown of glucose, also displays latitudinal clinality, such that the allozyme with low activity is found at higher frequencies at higher latitudes (Bubliy et al., 1999; Oakeshott et al., 1983; Singh et al., 1982). We can speculate that this may contributie to the higher levels of free glucose, however functional tests remain an important next step. Overall, this suggests that geographic and environmental gradients are among the selective forces that mediate genetic variation in glucose metabolism and its impact on behavior.

Clinal patterns have been found for numerous phenotypes and their underlying genetic architecture, including allozyme variants, sequence variants, and chromosome inversions (Fabian et al., 2012; Knibb, 1982; Kolaczkowski et al., 2011; Oakeshott et al., 1983; Sezgin et al., 2004), suggesting that convergent evolution in similar environments may have shaped variation in these traits, albeit in different parts of the world. The expansion to northern habitats is suggested to be a relatively recent phenomenon in the history of *D. melanogaster.* This species is thought to have migrated from equatorial zones to northern latitudes 10,000-20,000 years ago (Begun and Aquadro, 1993; David and Capy, 1988). Migration to certain geographic areas, such as North America and Australia, are thought to be much more recent, and as late as the last 200 years (Hoffmann and Weeks, 2007; Keller, 2007; Knibb, 1982). It is possible that reduced sleep is associated with this migration and is an evolutionary adaptation to seasonal changes in temperature and light cycle (Adrion et al., 2015; Li and Stephan, 2006). Achieving a better understanding of the evolution and biogeography of *D. melanogaster* in the geographic regions investigated in this study will help inform the relationship between sleep and the metabolic traits measured here.

Overall, our results reveal dramatic variation in sleep and metabolic regulation. The observations associating clinal variation with naturally occurring differences in sleep duration suggests that environmental gradients are potent selective forces that can shape behavior. We demonstrate here that latitude and average temperature contribute to variation in sleep-related behaviors, but likely there are numerous additional environmental factors that modulate sleep and metabolic function.

## ACKNOWLEDGEMENTS

We would like to thank Masato Yoshizawa (U. Hawai’i) and Hersh Chaitin (FAU) for their helpful discussions on statistical analyses. We would also like to thank members of the Keene lab for technical support. This work was supported by the National Institutes of Health award R01NS085152 to A.C.K.

## AUTHOR CONTRIBUTIONS

A.C.K. and E.B.B. designed the study and wrote the manuscript. E.B.B., J.T., V.R., and A.K. conducted the sleep and starvation experiments. R.A.B. and J.R.D. performed the nutrient storage measurements. E.B.B. analyzed the data. All authors read and approved of the manuscript.

